# The crystal structure of heme *d_1_* biosynthesis-associated small c-type cytochrome NirC reveals mixed oligomeric states *in crystallo*

**DOI:** 10.1101/2020.02.21.959056

**Authors:** Thomas Klünemann, Steffi Henke, Wulf Blankenfeldt

## Abstract

Monoheme c-type cytochromes are important electron transporters in all domains of life. They possess a common fold hallmarked by three α-helices that surround a covalently attached heme. An intriguing feature of many monoheme c-type cytochromes is their capacity to form oligomers by exchanging at least one of their α-helices, which is often referred to as 3D domain swapping. Here, we have determined the crystal structure of NirC, a c-type cytochrome co-encoded with other proteins involved in nitrite reduction by the opportunistic pathogen *Pseudomonas aeruginosa*. Crystals diffracted anisotropically to a maximum resolution of 2.12 Å (spherical resolution 2.83 Å) and initial phases were obtained by Fe-SAD phasing, revealing the presence of eleven NirC chains in the asymmetric unit. Surprisingly, these protomers arrange into one monomer and two different types of 3D-domain-swapped dimers, one showing pronounced asymmetry. While the simultaneous observation of monomers and dimers probably reflects the interplay between high protein concentration required for crystallization and the structural plasticity of monoheme c-type cytochromes, the identification of conserved structural motifs in the monomer together with a comparison to similar proteins may offer new leads to unravel the unknown function of NirC.

**Synopsis:** The crystal structure of the c-type cytochrome NirC from *Pseudomonas aeruginosa* has been determined and reveals the simultaneous presence of monomers and 3D-domain-swapped dimers in the same asymmetric unit.

## 1. Introduction

Monoheme c-type cytochromes are a subfamily of c-type cytochrome proteins and well-known electron transporters in the respiratory chain of organisms in all domains of life. In bacteria, they are also involved in H_2_O_2_-scavenging and work as electron entry point for the nitrate-, nitrite-, and nitric oxide reductases, which function as terminal oxidases in the respiratory chain under anaerobic conditions (denitrification) (Bertini *et al*., 2006). The monoheme c-type cytochrome domain features three α-helices that surround one c-type heme covalently attached to the cysteines of the characteristic sequence motif CXXCH *via* thioether bonds. The central iron cation of heme is usually coordinated by a histidine residue at its proximal site, whereas a histidine or, more rarely, a methionine ligates the other side in most cases. Many monoheme c-type cytochromes can form dimers and oligomers by exchanging the N- or C-terminal α-helix, which offers attractive protein engineering opportunities (Nagao *et al*., 2015; Hayashi *et al*., 2012; Hirota *et al*., 2010). We have recently embarked on a structural biology program to investigate enzymes involved in the biosynthesis of heme *d_1_*, an essential cofactor of the *cd_1_* nitrite reductase NirS, which catalyses the reduction of nitrite to nitric oxide in e.g. the opportunistic pathogen *Pseudomonas aeruginosa* (Schobert & Jahn, 2010). The *nir*-operon of *P. aeruginosa*, which encodes most of the enzymes involved in heme *d_1_* biosynthesis in addition to *nirS* itself, also contains open reading frames for two monoheme c-type cytochromes termed *nirM* and *nirC* (Figure 1) (Arai *et al*., 1990; Kawasaki *et al*., 1995). Whereas NirM is absent in many NirS-producing bacteria and has been assigned to function as an electron transporter for the nitrite reductase (Hasegawa *et al*., 2001), orthologs of NirC are found in nearly all NirS producers, but its function is subject to ongoing discussions. It has been demonstrated that NirC of *Pseudomonas aeruginosa* can act as an electron donor for the nitrite reductase *in vitro* (Hasegawa *et al*., 2001). However, the redox potential of NirC from *Paracoccus pantothrophus* has been found to not be optimal for the transport of electrons from the bc1-complex to nitrite reductase (Zajicek *et al*., 2009; Hasegawa *et al*., 2001). The double knockout of the two better-suited electron transporters cytochrome *c*550 or pseudoazurin rendered *Paracoccus denitrificans* unable to denitrify despite the presence of NirC (Pearson *et al*., 2003). Further, mutants of *Paracoccus denitrificans* and *Pseudomonas fluorescens* impaired in their ability to express *nirC* are unable to produce active nitrite reductase due to the lack of heme *d_1_*. Together, these findings implicate the role of NirC to lie in heme *d_1_* biosynthesis (Boer *et al*., 1994; Ye *et al*., 1992), which could, however, not be confirmed in *P. aeruginosa* (Hasegawa *et al*., 2001). To further investigate NirC, we have determined the crystal structure of heterologously produced *P. aeruginosa* NirC. We find a unique crystal packing that contains monomeric NirC together with two different dimeric forms. To our knowledge, this is the first time that monomers and dimers of a monoheme c-type cytochrome have simultaneously been observed in the same asymmetric unit.

**Figure 1.**
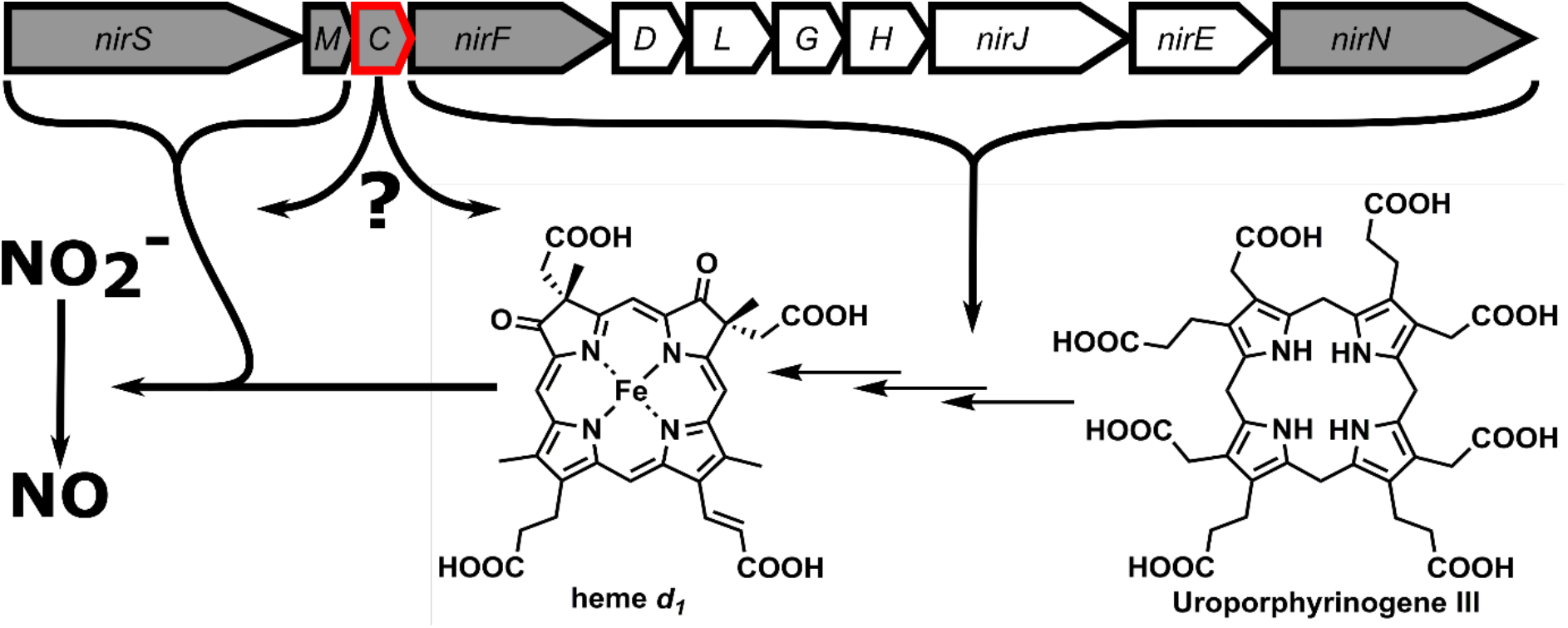
Depiction of the *nir*-operon indicating the function of each gene product in nitrite reduction and heme *d_1_* biosynthesis. A grey background highlights the periplasmic location of the gene product.

## 2. Materials and methods

### 2.1. Macromolecule production

The gene encoding NirC was amplified by polymerase chain reaction from genomic DNA extracted from *P. aeruginosa* PA14, omitting the signal peptide for periplasmic export as determined by SignalP 5.0 (Almagro Armenteros *et al*., 2019). The fragment was cloned between the restriction sites NotI/KpnI of a modified pCOLA-Duet plasmid (pVP008-ompA), yielding an expression vector that produces NirC with an N-terminal ompA signal peptide for periplasmic export in *E. coli* followed by a Tobacco Etch Virus (TEV) protease-cleavable StrepII-tag. Residue E71 was mutated to alanine by QuikChange Mutagenesis for reasons outlined below. The success of cloning and mutagenesis was verified by the Eurofins Genomic sequencing service.

*E. coli* C43(DE3) (Miroux & Walker, 1996) was cotransformed with the NirC expression plasmid and with pEC86, a plasmid containing the cytochrome *c* maturation system of *E. coli*, which is necessary for the covalent attachment of heme to a suited polypetide chain (Arslan *et al*., 1998). LB medium (8 x 1 l) supplemented with 34 mg/L chloramphenicol, 30 mg/L kanamycin, 200 μM δ-aminolevulinic acid and 12.5 μM Fe(II)SO4 was inoculated with 10 mL overnight culture, cultivated at 37 °C with mild shaking until an optical density between 0.6 and 0.8 was reached. Cultures were then induced with 1 mM isopropyl β-D-1-thiogalactopyranoside and further incubated at 20 °C for 20 h. Cells were harvested by centrifugation and pellets were stored at −20 °C until needed.

Cell pellets were thawed, resuspended in lysis buffer (50 mM Tris-HCl pH 8.0, 150 mM NaCl) supplemented with protease inhibitors (cOmplete mini EDTA-free) and lysed by sonication. The lysate was centrifuged 30 min at 36,000 *g* followed by an additional centrifugation of the supernatant for 60 min at 100,000 *g* to remove cell debris.

The cleared cell lysate was loaded on a StrepTactin-HC column (IBA, Göttingen, Germany) equilibrated in protein buffer (10 mM Tris-HCl pH 8.0, 150 mM NaCl) and connected to an Äkta Purifier FPLC system (GE Healthcare, Boston, U.S.A.). The column was washed with five column volumes and subsequently eluted with three column volumes of protein buffer containing 5 mM d-desthiobiotin. Red colouration of elution fractions and UV-Vis absorption at 410 nm confirmed the presence of heme. Fractions containing no impurities as judged by SDS-PAGE analysis were pooled and digested with TEV-protease at 4 °C overnight with simultaneous dialysis against protein buffer. To remove undigested protein, the resulting solution was again loaded onto the StrepTactin column and the flowthrough was collected. As final purification step, NirC was subjected to size exclusion chromatography using a HiLoad 16/60 Superdex 75 column (GE Healthcare, Boston, U.S.A.). Pure protein was concentrated to 40 mg/mL with Vivaspin 20 (MWCO 3000) ultrafiltration units (Sartorius AG, Göttingen, Germany) and concentration was determined photometrically, using a NanoDrop2000 (ThermoFisher Scientific, Waltham, U.S.A.) and a calculated extinction coefficients of ε_280_=11 mM^-1^*cm^-1^.

**Table 1.**
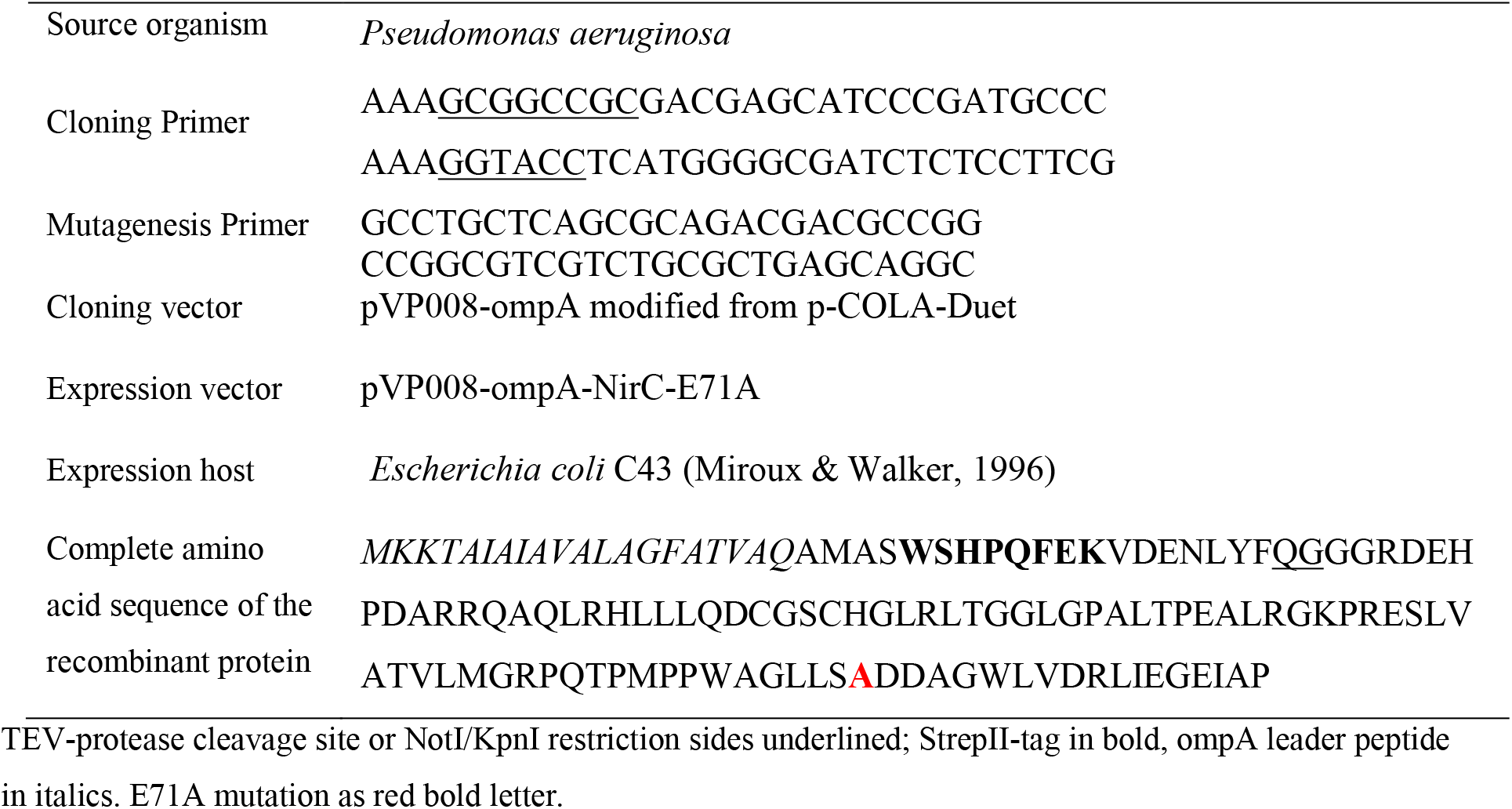
Macromolecule production information

### 2.2. Crystallization

Crystallization experiments were set up using a HoneyBee 961 pipetting robot (Digilab Genomic Solutions, Hopkinton, U.S.A.) and monitored with a Rock Imager 1000 automated microscope (Formulatrix, Bedford, U.S.A.). Optimization screens were prepared utilizing a Formulator pipetting robot (Formulatrix, Bedford, U.S.A.). Experiments to crystallize wild type NirC never yielded well-diffracting crystals. Therefore, a mutation (E71A) as suggested by the surface entropy reduction server (Goldschmidt *et al*., 2007) was introduced, leading to crystal growth in a few conditions of the Cryos sparse matrix screen (Qiagen, Hilden, Germany). The respective precipitants were further optimized by varying the concentration of precipitant and salts as well as the pH. Crystals usually began to appear after one week and took three more weeks to grow to full size (50-100 μm) (Supplementary Figure 1). Crystals were harvested and directly flash-cooled in liquid nitrogen without the addition of cryoprotectants.

**Table 2.** Crystallization

|  | Crystal used for SAD phasing | Crystal used for final structure refinement |
| --- | --- | --- |
| Method | Sitting drop vapour diffusion | Sitting drop vapour diffusion |
| Plate type | Intelli 96-3 | Intelli 96-3 |
| Temperature (K) | 293 | 293 |
| Protein concentration (mg/ml) | 27 | 40 |
| Buffer composition of protein solution | 10 mM Tris-HCl pH 8.0<br>150 mM NaCl | 10 mM Tris-HCl pH 8.0, 150 mM NaCl |
| Composition of reservoir solution | 15% glycerol<br>8.5 mM $\text{NiCl}_2$<br>85 mM Tris-HCl pH 8.5<br>17% PEG-MME 2000 | 15.33% glycerol<br>0.1 M MES-NaOH pH 6.3<br>10.8% PEG 20000 |
| Volume and ratio of drop | 200 nL : 200 nL | 200 nL : 200 nL |
| Volume of reservoir | 60 $\mu\text{L}$ | 60 $\mu\text{L}$ |

### 2.3. Data collection and processing

Data were collected at beamline P11 of the PETRAIII synchrotron (DESY, Hamburg, Germany) (Burkhardt *et al*., 2016). The anomalous data set was processed using DIALS (Winter *et al*., 2018), POINTLESS (Evans, 2011) and AIMLESS (Evans & Murshudov, 2013) of the CCP4 suite (Winn *et al*., 2011). Despite low completeness of the highest resolution shell the data set was sufficient to obtain initial phases via SAD phasing. The data set used for final refinement was processed using autoPROC (Vonrhein *et al*., 2011) executing XDS (Kabsch, 2010), POINTLESS (Evans, 2011), AIMLESS (Evans & Murshudov, 2013) and STARANISO (Tickle *et al*., 2018). Anisotropic data truncation was applied assuming a local *I*/σ(*I*) of 1.2 to include useable data beyond the spherical resolution of 2.83 Å. The extent of anisotropy is visualised in Supplementary Figure 5.

**Table 3.** Data collection and processing Values for the highest resolution shell are given in parentheses.

|  | Anomalous Dataset | Native dataset |
| --- | --- | --- |
| Diffraction source | Petra III beamline P11 | Petra III beamline P11 |
| Wavelength (Å) | 1.738 | 1.033 |
| Temperature (K) | 100K | 100K |
| Detector | Pilatus 6M | Pilatus 6M |
| Crystal-detector distance (mm) | 544 | 457.5 |
| Rotation range per image (°) | 0.1 | 0.1 |
| Total rotation range (°) | 1080 | 360 |
| Exposure time per image (s) | 0.1 | 0.04 |
| Space group | $P2_12_12_1$ | $P2_12_12_1$ |
| $a, b, c$ (Å) | 77.7, 80.6, 200.0 | 76.8, 80.2, 195.7 |
| $\alpha, \beta, \gamma$ (°) | 90, 90, 90 | 90, 90, 90 |
| Resolution range (Å) | 200-3.4 (3.7-3.4) | spherical: 99.2-2.8 (2.9-2.8)<br>ellipsoidal: 99.16-2.19 (2.47-2.19) |
| Diffraction limit along principal axes of fitted ellipsoid | - | 2.98 ($a^*$)<br>2.91 ($b^*$)<br>2.12 ($c^*$) |
| Total No. of reflections | 368556 (6018) | 460561 (18799) |
| No. of unique reflections | 13722 (873) | 35413 (1772) |
| Completeness spherical (%) | 77.3 (24.9) | 53.6 (8.9) |
| Completeness ellipsoidal (%) | - | 92.7 (74.8) |
| Multiplicity | 26.9 (6.9) | 13.0 (10.6) |
| $\langle I/\sigma(I) \rangle$ | 10.5 (2.2) | 10.9 (1.8) |
| CC<sub>1/2</sub> | 0.993 (0.721) | 0.998 (0.610) |
| R<sub>p.i.m.</sub> | 0.046 (0.404) | 0.058 (0.416) |
| Overall *B* factor from Wilson plot (Å<sup>2</sup>) | 63 | 23 |

### 2.4. Structure solution and refinement

The structure was solved by SAD phasing with anomalous differences caused by the heme iron atoms, using the phasing pipeline Crank2 (Skubák & Pannu, 2013) executing ShelxC/D (Sheldrick, 2008) for substructure determination, Parrot (Cowtan, 2010) for density modification and Buccaneer (Cowtan, 2006) for model building. The iron substructure determination showed an occupancy drop for the 12^th^ atom and beyond, hinting at the presence of eleven chains in the asymmetric unit. Automated model building produced a model consisting of 940 amino acids and having an R_work_ of 35%. Inspection of the electron density showed that not all of the eleven chains were built correctly, requiring manual adjustments in Coot (Emsley & Cowtan, 2004) and further refinement in REFMAC5 (Murshudov *et al*., 2011) with enabled NCS restraints. Final refinement was performed against the higher resolution native data set, using phenix.refine (Afonine *et al*., 2012) while lifting the applied NCS restrains, adding water molecules, riding hydrogen atoms and applying TLS refinement. Depictions of the structure were made with PyMol (Schrödinger, 2015).

**Table 4.**
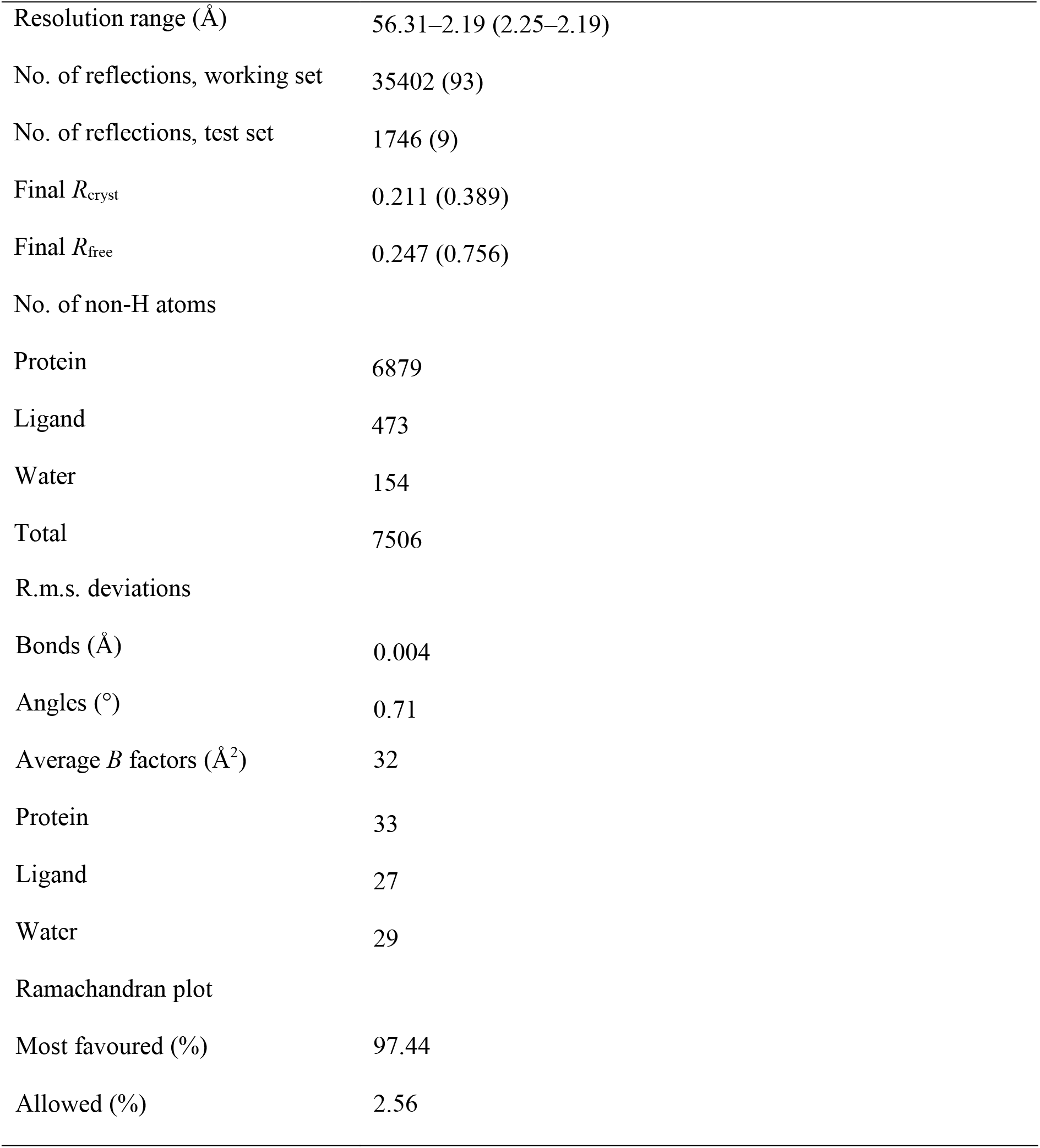
Structure solution and refinement Values for the highest resolution shell are given in parentheses.

|  |  |
| --- | --- |
| Resolution range (Å) | 56.31–2.19 (2.25–2.19) |
| No. of reflections, working set | 35402 (93) |
| No. of reflections, test set | 1746 (9) |
| Final *R*<sub>cryst</sub> | 0.211 (0.389) |
| Final *R*<sub>free</sub> | 0.247 (0.756) |
| No. of non-H atoms |  |
| Protein | 6879 |
| Ligand | 473 |
| Water | 154 |
| Total | 7506 |
| R.m.s. deviations |  |
| Bonds (Å) | 0.004 |
| Angles (°) | 0.71 |
| Average *B* factors (Å<sup>2</sup>) | 32 |
| Protein | 33 |
| Ligand | 27 |
| Water | 29 |
| Ramachandran plot |  |
| Most favoured (%) | 97.44 |
| Allowed (%) | 2.56 |

### 2.5. Size exclusion chromatography coupled to a multi-angle laser light detector

To assess the molecular weight of NirC-E75A the isolated and concentrated protein solution (40 mg/mL) was subjected to SEC-MALS experiments utilizing an Agilent Technologies 1260 Infinity II HPLC system (Santa Clara, U.S.A.) equipped with a Wyatt Optilab rEX diffraction index detector and a Treos II multi angle laser light scattering detector (Wyatt, Santa Barbara, U.S.A.). Separation was achieved with a Superdex 75 Increase 10/300 GL column (GE Healthcare, Boston, U.S.A.) with protein buffer as eluent. For calculation of the molecular mass, a dn/dc of 0.195 mL/g was assumed.

### 2.6. Structural Bioinformatics

Functional orthologs of NirC were identified by extracting orthologs from the OMA database (Altenhoff *et al*., 2018) and followed by manually filtering for the presence of the genes encoding nitrite reductase NirS and heme *d_1_* biosynthesis enzyme NirF in their genetic neighbourhood. After aligning the functional orthologs with ClustalΩ (Ashkenazy *et al*., 2016), sequence conservation was mapped onto the NirC monomer structure utilizing the ConSurf server (Figure 3 and Supplementary Figure 2) (Sievers *et al*., 2011). Structural homologs were identified with the DALI server (Holm & Laakso, 2016).

**Figure 2.**
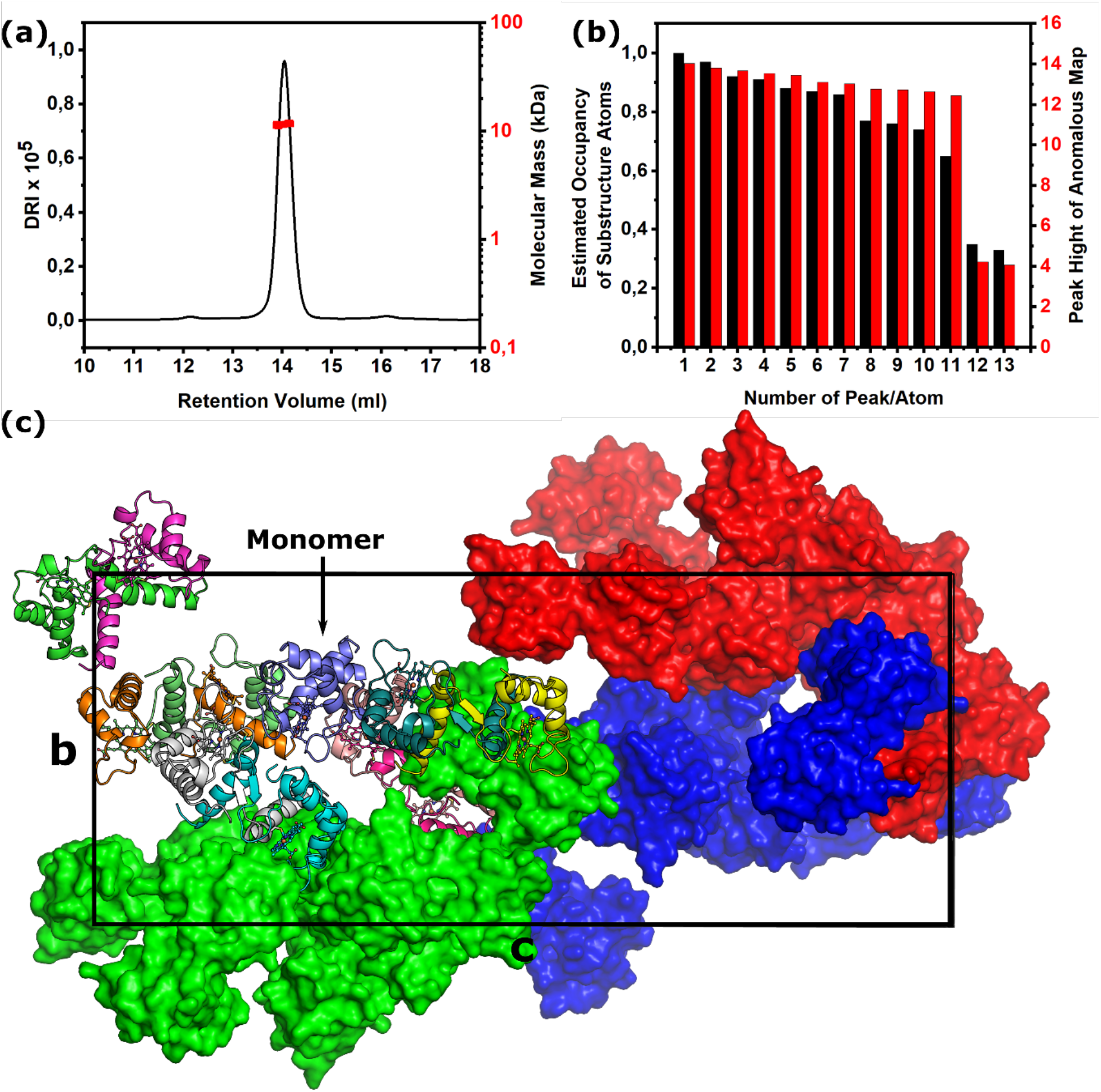
Panel (a) shows the SEC-MALS chromatogram. Bar diagrams in (b) show the occupancy of atoms of the initially determined substructure as calculated with ShelxD (Sheldrick, 2008) and the height of the anomalous map peak calculated with phases of the final structure by ANODE (Thorn & Sheldrick, 2011). The drop after the 11^th^ site indicates that only eleven iron atoms are present. Panel (c) depicts the unit cell. One asymmetric unit is shown as in cartoon representation of the peptide backbone with the covalently attached heme as ball-and-stick model and each chain coloured individually. The other asymmetric units are depicted as surface model coloured in red, blue and green.

**Figure 3.**
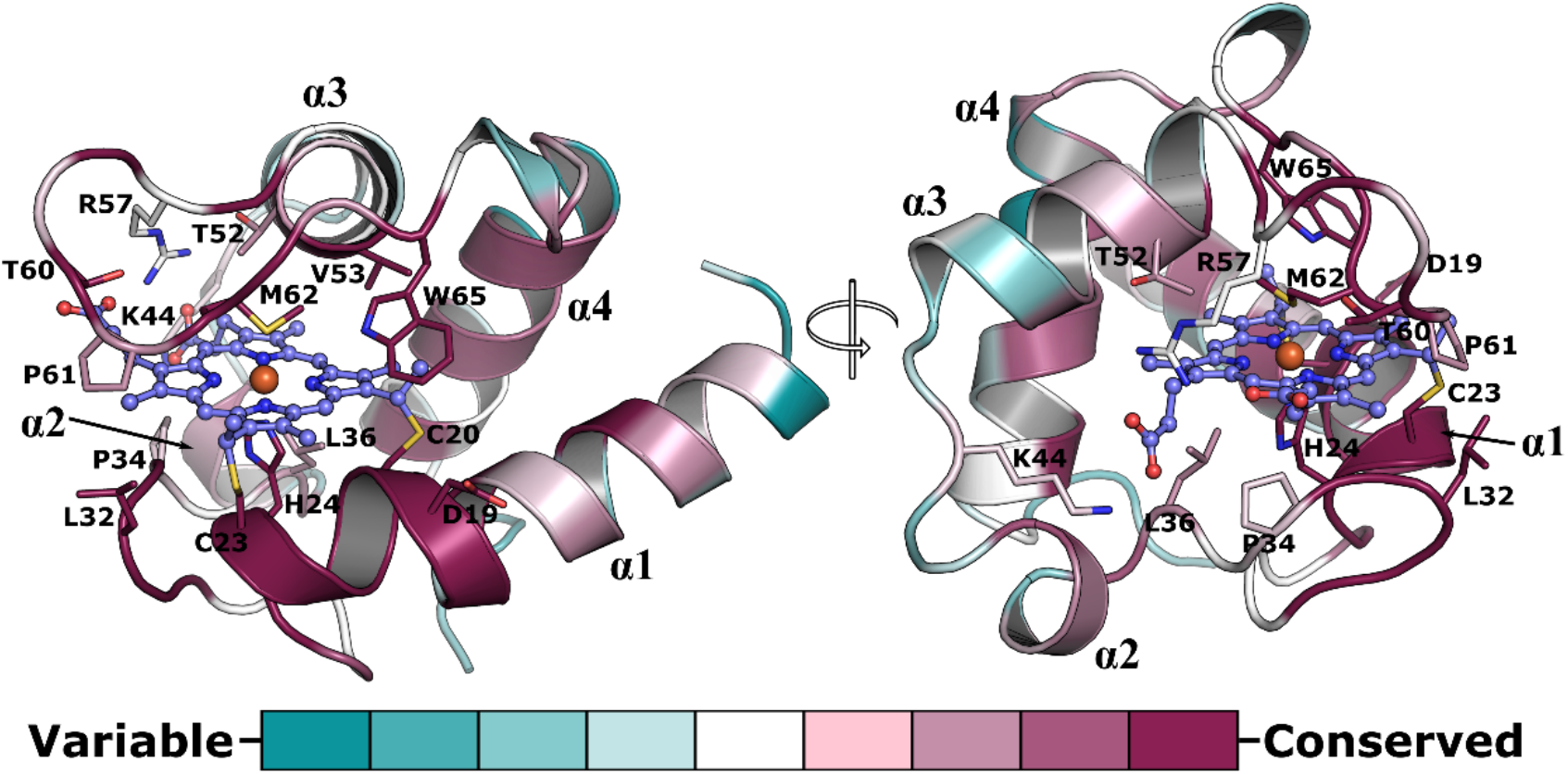
The NirC monomer shown as a cartoon with residues at a maximum distance of 4 Å to the covalently attached heme presented as sticks. Colours are based on sequence conservation as defined by Consurf (Ashkenazy *et al*., 2016). The sequence alignment used for determining conservation is shown in Supplementary Figure 3.

## 3. Results and discussion

### 3.1. Simultaneous presence of NirC monomers and dimers in the asymmetric unit

Approximately 2 mg of pure heme-bound NirC per litre of *E. coli* culture could reliably be obtained with the procedure described above. Size exclusion chromatography as well as multi-angle light scattering suggested that the concentrated protein solution used in crystallisation trials only contains monomers with a molecular mass of 11.5 kDa ± 0.9% (10.5 kDa expected, Figure 2). Because the wildtype protein failed to crystallize, we attempted to reduce the surface entropy by mutating E71 to alanine, finally yielding cube-shaped crystals that diffracted anisotropically and possessed a very large orthorhombic unit cell (Supplementary Figure 1). To obtain initial phases, we made use of the presence of anomalous differences in diffraction data collected at the iron K-edge, leading to the positioning of eleven iron atoms. By assuming that NirC contains one iron atom bound to the covalently attached heme, this suggests a solvent content of 58%. A combination of automated model building and manual adjustments led to a final model with R-factors R_work_ = 21.1% and Rfree = 24.7%, indeed consisting of eleven chains in the asymmetric unit. Interestingly, despite using monomeric NirC for crystallization, ten of these chains form dimers by exchanging their N-terminal α-helices recognizable by different traces in the 2Fo-Fc density for the “hinge loop” (G25-G33; Figure 5). The dimers fall into two different types as outlined below. Surprisingly, the eleventh chain is monomeric, i.e. the crystal form obtained here contains both dimers and monomers of NirC at the same time. Analysis of the crystal packing reveals that the surface entropy reducing mutation of E71A enables packing interactions that are not possible with the glutamate sidechain. An example is shown in Supplementary Figure 6. Importantly, the mutation is not part of the dimer interfaces or the hinge loop, suggesting that it does not cause the simultaneous presence of different NirC oligomeric states observed here.

**Figure 4.**
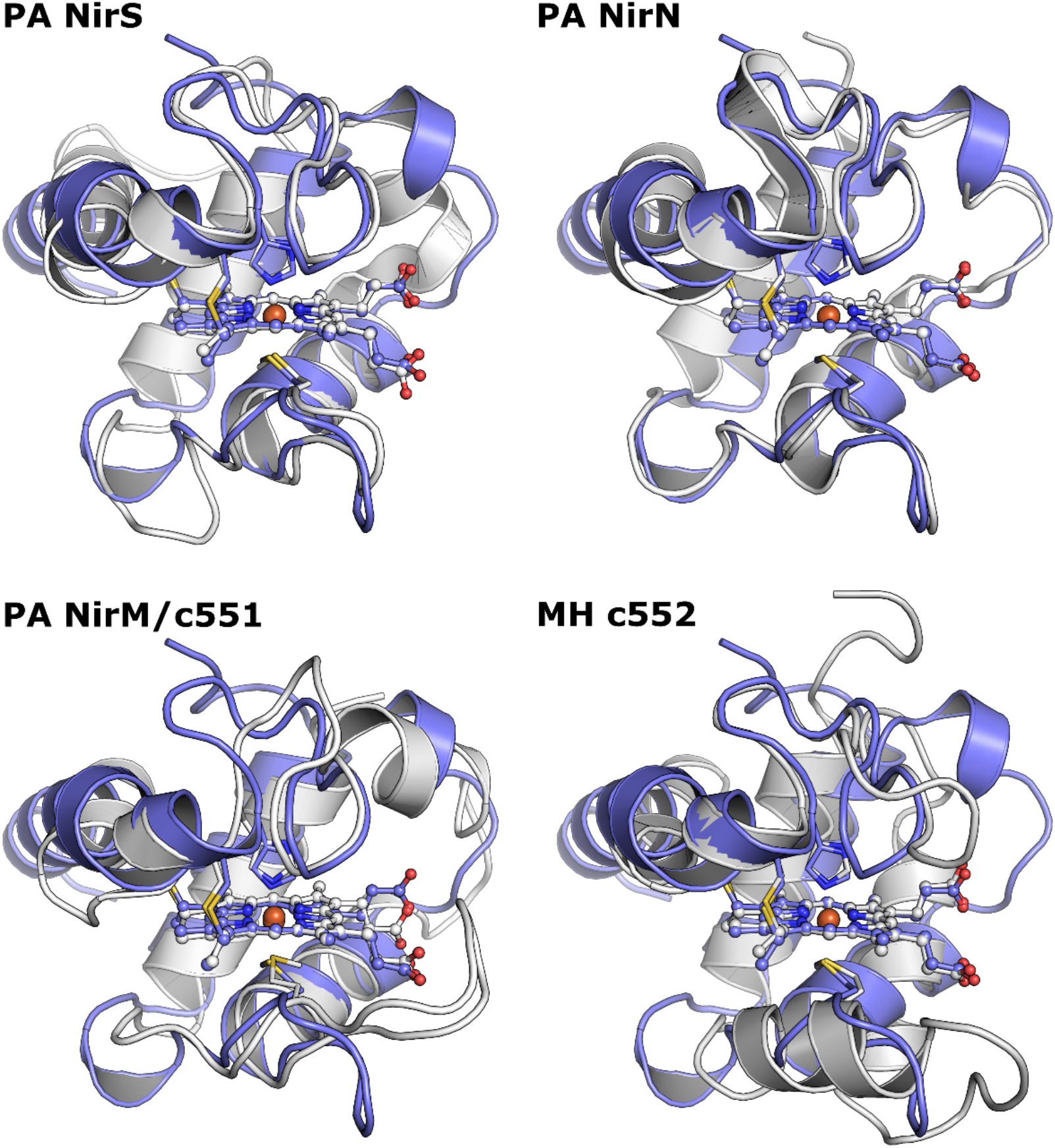
Comparison of the structure of monomeric NirC with the cytochrome *c* domains of NirS and NirN as well as with NirM from *Pseudomonas aeruginosa* or *cyt c552* of *Marinobacter hydrocarbonoclasticus* after superposition of the covalently attached heme. All structures are depicted as cartoons and the covalently attached heme as a ball-and-stick model. NirC is shown in blue and the compared structure in white.

**Figure 5.**
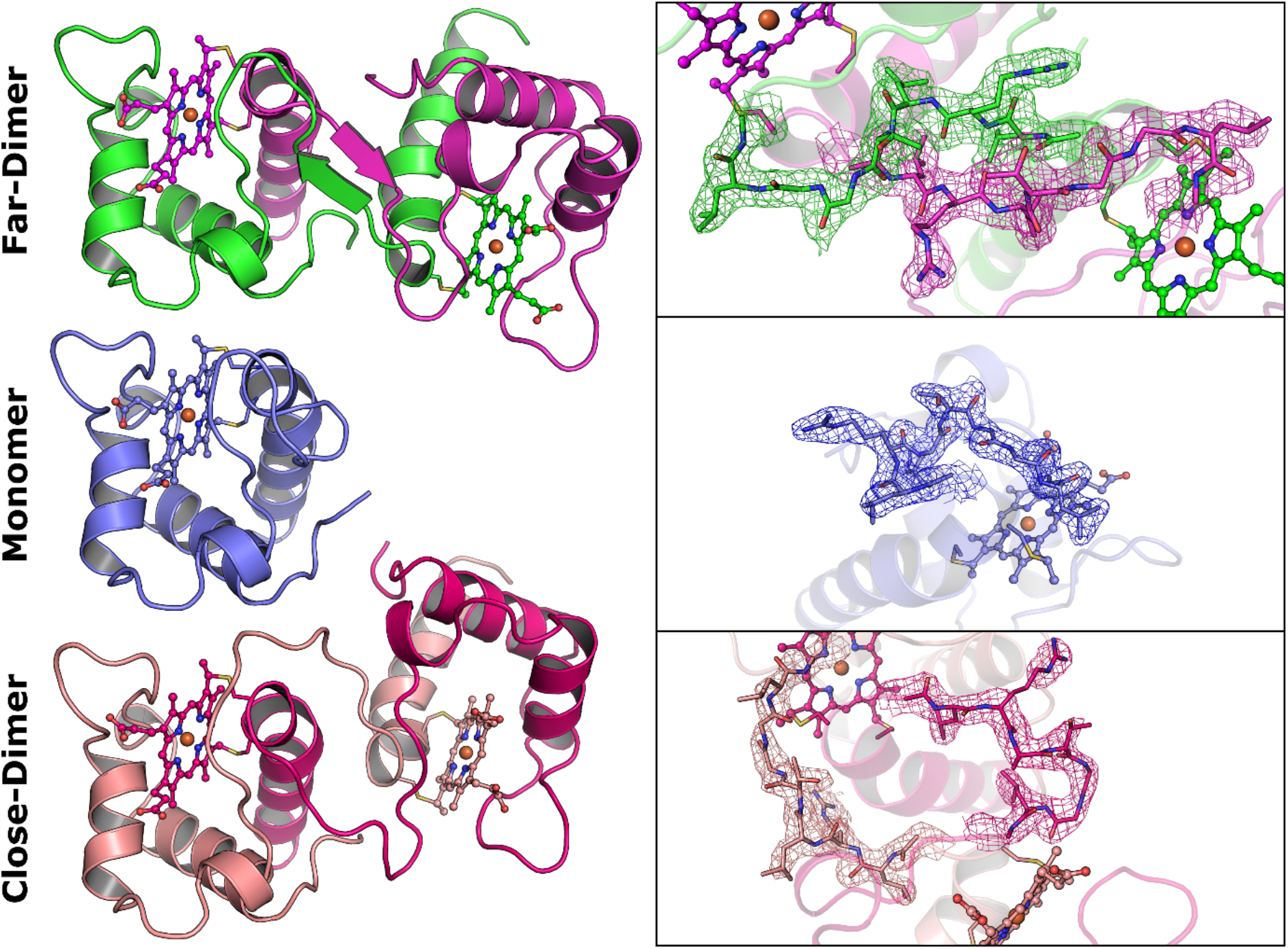
Depiction of the two different oligomerization states found in the crystal structure of NirC. On the left the NirC monomer and both dimer conformations are shown as described in Figure 1 after superposition of one protomer. The backbone of the NirC is shown as a cartoon with the covalently attached heme as ball-and-stick model. On the right, residues of the hinge loop are depicted as stick models with a 2FoFc map at a σ-level of one, coloured according to the associated chain and individually orientated to allow unobstructed view.

The formation of dimers from monomeric c-type cytochromes is a well-known phenomenon (Hirota, 2019). It has been shown that the folding of horse cytochrome *c* starts with hydrophobic interactions between the N- and C-terminal helix, yielding 3D-domain-swapped oligomers if the helices belong to different chains (Parui *et al*., 2013). Therefore, denaturation and subsequent refolding at high protein concentration is often utilized to produce such dimers (Nagao *et al*., 2015; Hayashi *et al*., 2012). Apparently, the high NirC concentration and the presence of precipitants in the crystallization experiments performed here is sufficient to allow an equilibrium between dimers and monomers.

Whereas 3D domain swapping is often observed in crystal structures (reviewed in (Liu & Eisenberg, 2002)), the simultaneous presence of different homooligomeric states of one protein in the same asymmetric unit is extremely rare, and we are aware of only two other unambiguous examples, namely the FeSII protein from *Azotobacter vinelandii* (PDB: 5FRT; unpublished) and the engineered tenascin protein (PDB: 2RBL) (Hu *et al*., 2007). In both cases, the asymmetric unit was found to contain monomers and 3D-domain-swapped dimers, similar to the crystal structure of NirC reported here. In addition, there may be less obvious instances that do not involve 3D domain swapping, such as a non-oligomerizing variant of the auxin response factor from *Arabidopsis thaliana* (PDB: 4NJ7). This protein crystallized with 16 chains in the asymmetric unit, which may be regarded as dimers and pentamers (Korasick *et al*., 2014), although it also seems possible to explain the packing by assuming only monomers.

### 3.2. NirC shows high similarity to other monoheme c-type cytochrome domains of the *nir*-operon

NirC displays a typical monoheme c-type cytochrome fold containing of three α-helices (α1, α3, α4) that surround a thioether-bound heme *c* moiety (Figure 3). The iron cation is coordinated by H24 and M62, which is consistent with mutagenesis studies performed with the orthologue from *Paracoccus panthotrophus* (Zajicek *et al*., 2009). The heme-binding crevice consists of a predominantly hydrophobic pocket established by L32, P34, L36, T52, V53, T60, P61 and W65, which are also highly conserved in NirC proteins from other species. In *P. aeruginosa* NirC, the propionate groups of heme form H-bonds with K44 and R57, two residues that are replaced with hydrophobic amino acids in some species, probably reflecting the fact that the heme propionate groups are solvent-exposed and do not require stabilizing interactions with the protein to be accommodated. Sequence alignment reveals that in NirC the heme attachment motif (CXXCH) is always led by a conserved aspartate (D19), whose sidechain is solvent exposed, and followed by a conserved GGLG motif, residing on a loop connecting helix α2 and α3 (Supplementary Figure 3). Together with residues in the neighbourhood of the iron-ligating M62, these motifs create a conserved surface patch, which may be required to form an interaction site for other proteins such as e.g. the heme *d_1_* biosynthesis protein NirF, whose function is not clear but may lie in an oxidoreduction within the last steps of the pathway (unpublished, preprint version under https://doi.org/10.1101/2020.01.13.904656). Alternative hypotheses suggest an involvement as electron transporter for nitrite reductase NirS (Hasegawa *et al*., 2001). A role in heme *d_1_* biosynthesis itself, on the other hand, seems to be supported by the finding that similarity searches with DALI (Holm & Laakso, 2016) identify the cytochrome *c* domains of the *nir*-operon proteins nitrite reductase NirS and heme *d_1_* biosynthesis enzyme NirN as most similar to NirC, while NirM and cyt c552, the known electron transporters that interact with NirS in nitrite reduction, differ to a larger extent (Figure 4 and Supporting Table S1). In NirN and NirS, the cytochrome *c*-domains are covalently attached to a β-propeller domain and function as electron shuttles in the reactions catalysed by these enzymes. This suggests that NirC may perform a similar function with NirF, which is a standalone β-propeller protein with high similarity to the β-propeller domains of NirN and NirS. Interestingly, superposition of NirC and NirF onto the respective domains of NirS leads to a complex in which the conserved surface patch of NirC is oriented towards the propeller of NirF and L32 of the conserved GGLG motif is buried in a hydrophobic pocket (Supplementary Figure 4). This may further corroborate NirC’s involvement in heme *d_1_* biosynthesis.

### 3.3. NirC forms two different types of dimers

The dimers of NirC found here form by exchanging the N-terminal helix (P4-H24), which results in heme being coordinated by H24 and M62 coming from two different chains, similar to what has been observed for dimers of cyt c551 (NirM) from *P. aeruginosa* and cyt c552 from *Hydrogenobacter thermophiles* (HT) (Hayashi *et al*., 2012; Nagao *et al*., 2015). Notably, dimerization does not have a major impact on the structure of the heme binding site, indicating that monomers and dimers could display similar biochemical properties as was already found for other N-terminally domain-swapped cytochromes (Nagao *et al*., 2015; Hayashi *et al*., 2012).

Closer inspection reveals that the five dimers present in the asymmetric unit fall into two groups that are distinguishable by heme iron distance, internal symmetry and relative position of the hinge loop (25-GLRLTGGLG-33, Figure 5). In the three “far-dimers”, the average iron-to-iron distance is 28.5 ± 0.2 Å, whereas it is 24.9 ± 0.1 Å in the two “close-dimers”. Interestingly, the two dimeric forms of NirC display different amounts of internal symmetry, as can be quantified by the “GloA-score”. Naturally occurring homodimers are usually highly symmetric with GloA-scores under 0.4 Å, and scores above 3 Å are indicative of substantial asymmetry (Swapna *et al*., 2012). In the dimers of NirC observed here, the GloA-score is 0.9 ± 0.2 Å for the far-dimers, compared to 3.05 ± 0.01 Å for the close-dimers, indicative of substantial asymmetry in the latter, which is readily discernible after superposition of one promoter of a dimer onto the other (Supplementary Figure 2). The basis for the formation of different dimers in NirC seems to be rooted in the hinge loop, which with nine residues is much longer and allows more flexibility than the three residues found in PA cyt c551 or HT cyt c552 (Hayashi *et al*., 2012; Nagao *et al*., 2015). The importance of the length and hence the flexibility of the hinge loop is reflected in the observation that its elongation in HT cyt 552 induced different N- and C-terminally domain-swapped dimers (Ren *et al*., 2015). In NirC, the increased flexibility leads to anti-parallel β-sheet-like interactions of L26, R27 and L28 in the far-dimers and no interactions with elevated B-factors of the hinge loop in the close-dimers.

The relevance of dimer formation in NirC and the associated different levels of asymmetry are not clear at present. Previous work with eukaryotic monoheme c-type cytochromes suggested that 3D-domain-swapped oligomers are involved in apoptosis (Junedi *et al*., 2014), which seems to be in line with the finding that several other 3D-domain-swapped proteins have a regulatory function. Examples include bovine RNase, where only the dimer displayed toxicity towards tumour cells (Di Donato *et al*., 1995) or the regulatory domain of α-isopropylmalate synthase of *Mycobacterium tuberculosis*, which forms via 3D domain swapping (Koon *et al*., 2004). Because dimer formation of NirC is clearly linked to the hinge loop and because this loop his highly conserved amongst NirC homologues from different species, it is possible that dimers of NirC have a regulatory role as well. However, the fact that we did not detect dimers even in highly concentrated solutions together with the observation that crystallization took relatively long and required the introduction of a surface entropy reduction mutant that enabled crystal formation through new crystal contacts currently argues for a crystallization artefact that points towards the structural malleability of these proteins rather than a physiological importance of NirC dimers.

In summary, our work provides new insight into NirC, a monoheme c-type cytochrome likely involved in the biosynthesis of heme *d_1_*, the essential cofactor in nitrite reduction by denitrifying bacteria such as the opportunistic pathogen *Pseudomonas aeruginosa*. While the simultaneous observation of NirC monomers and dimers in the same asymmetric unit of the crystal form obtained here may reiterate typical properties of this protein family, the mapping of conserved sequence motifs onto the structure of the monomer may offer new avenues to decipher the function of NirC.

## Acknowledgements

We thank Dr. Stefan Schmelz for his support in interpreting SEC-MALS data and the beamline staff at P11 at the PETRAIII synchrotron (Deutsches Elektronensynchrotron DESY, Hamburg, Germany) for granting us excess to their facilities. Members of the ccp4 community are acknowledged for pointing us to other crystal structures with different oligomers in the same asymmetric unit. TK and SH were supported by the PROCOMPAS graduate school (grant GRK 2223/1 by the Deutsche Forschungsgemeinschaft).

## Supporting information

**Table S1.** Selected high scoring results of the Dali server discussed in this publication.

| Organism and<br>Name | PDB-ID | Z-score | lali | rmsd (Å) | %id | Citation |
| --- | --- | --- | --- | --- | --- | --- |
| PA NirN | 6rtd | 12.2 | 76 | 1.6 | 37 | (Klünemann<br>*et al.*, 2019) |
| PA NirS | 1nno | 12 | 81 | 2.3 | 26 | (Nurizzo *et al.*, 1998) |
| PA NirM/<br>cyt c551 | 351c | 7.8 | 71 | 2.5 | 25 | (Matsuura *et al.*, 1982) |
| MH cyt c552 | 1cno | 5.2 | 65 | 2.7 | 23 | (Brown *et al.*, 1999) |

**Figure S1.**
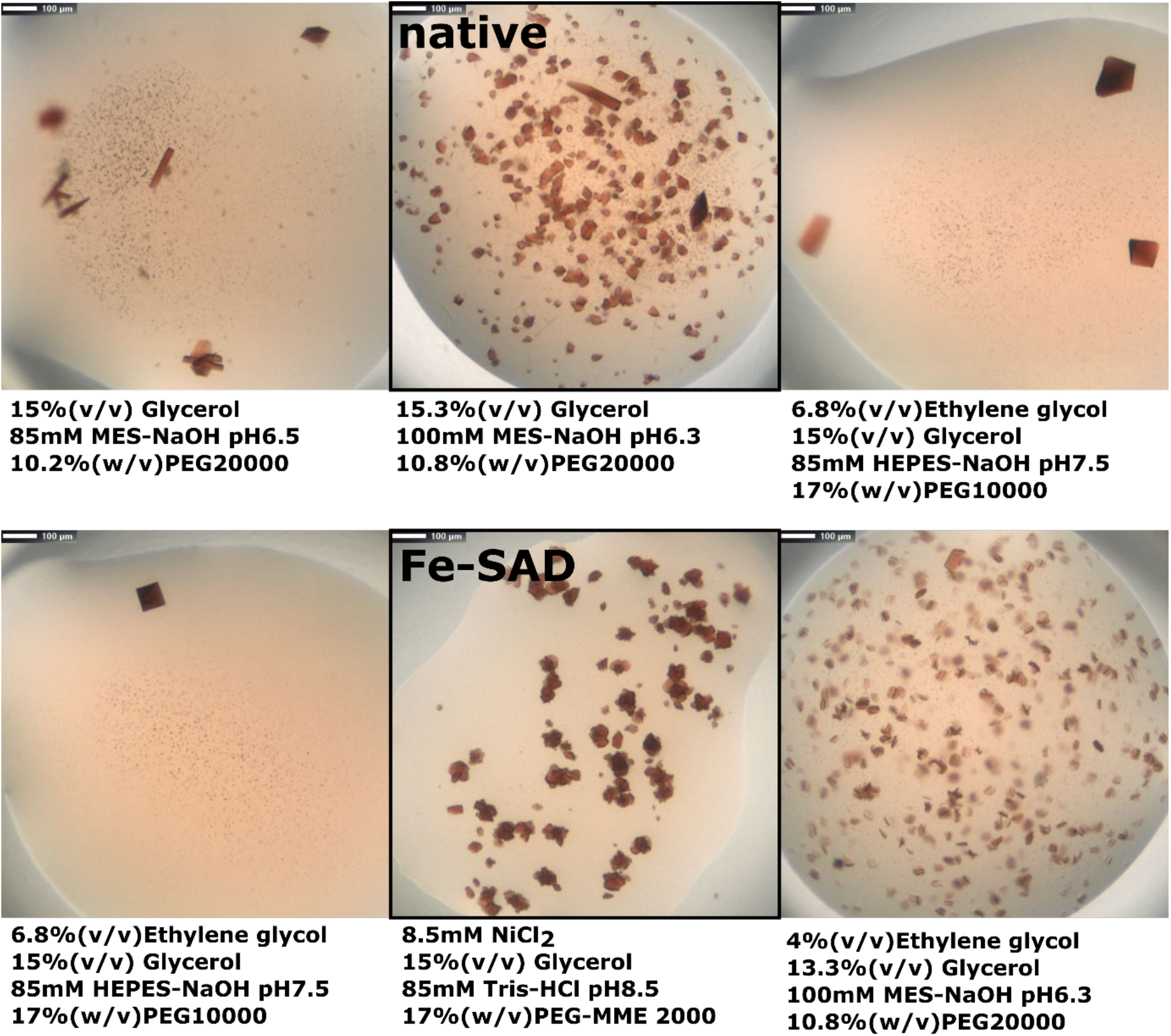
Crystals of NirC^E71A^ grown in different conditions used for diffraction experiments. Conditions used in this study are indicated by a black frame.

**Figure S2.**
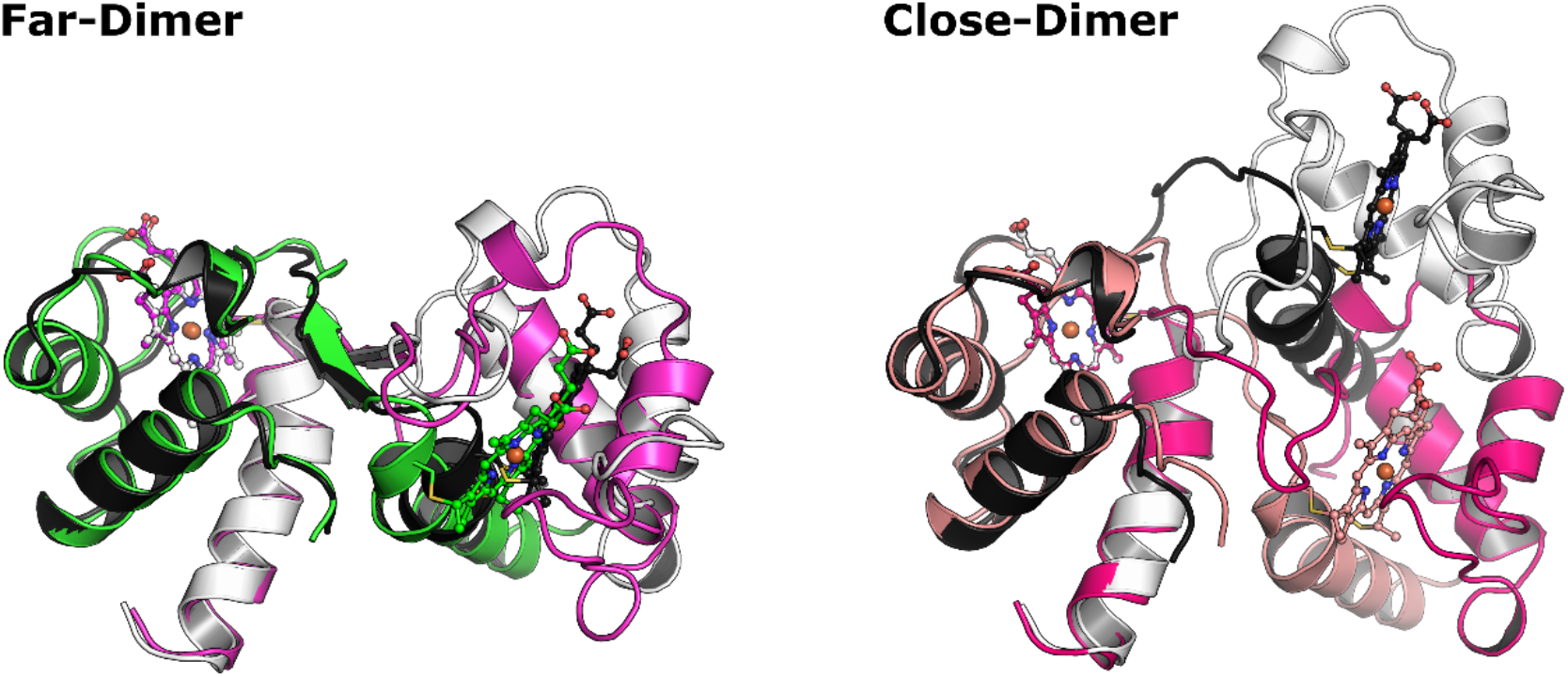
Depiction of NirC dimers. To highlight the asymmetry, one protomer was superposed with the copy of the opposite one. Except for the colouration of the copied dimer (black and white), the colour and representation of the model is the same as lied out in Figure 5 in the main text.

**Figure S3.**
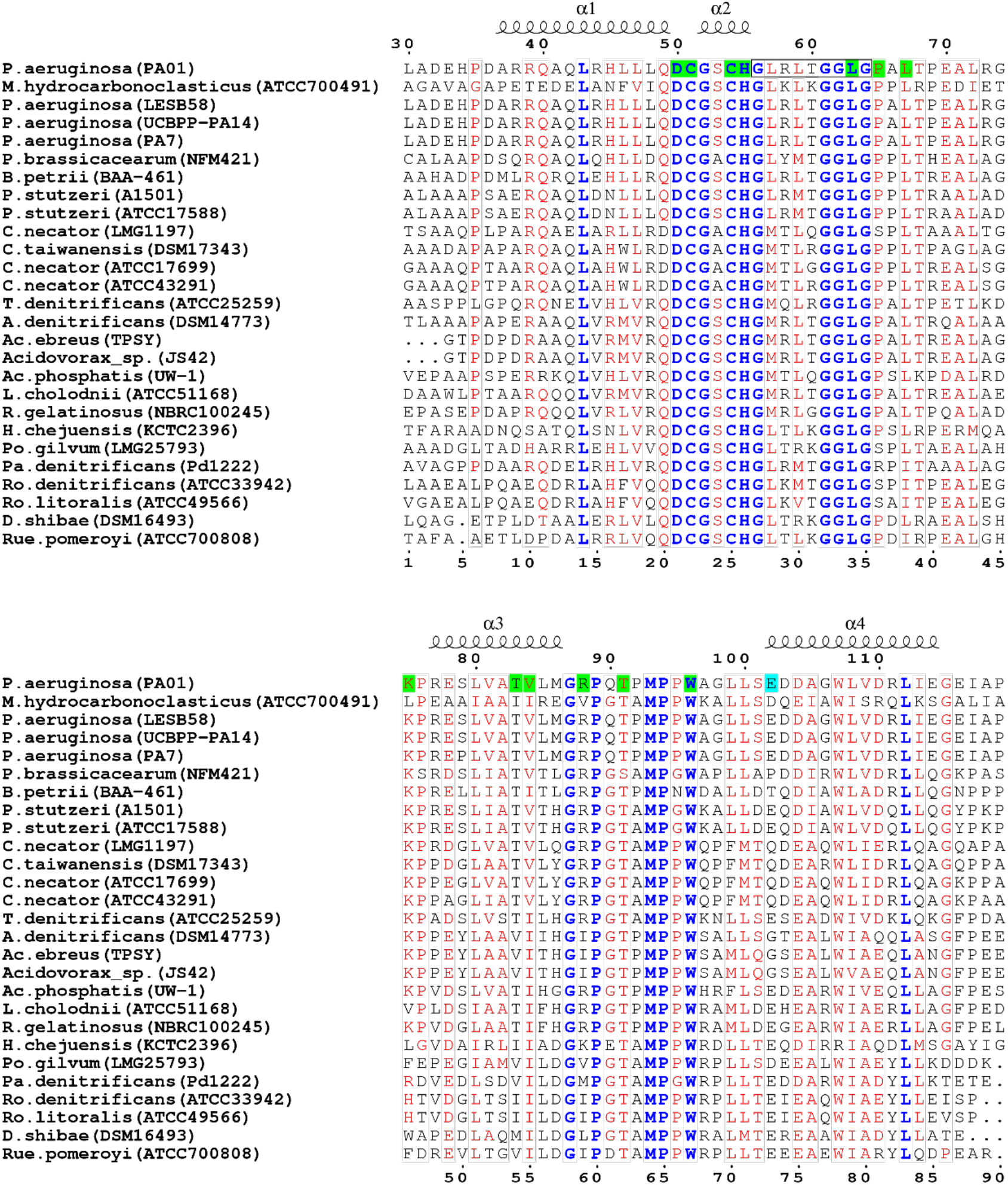
Sequence alignment of NirC orthologs from different denitrifying species extracted from the OMA server (Altenhoff *et al*., 2018). Green boxes mark heme-interacting residues. A cyan box indicates E71 mutated to alanine to enable crystal formation and the black box highlights residues belonging to the hinge loop, which adopts different conformation in the 3D domain swapped dimers. Bold blue letters indicate identity in all residues and red letters are equivalent to 70%. Sequence numbering is based on whole protein including the signal peptide. The figure was prepared with ESPript3.0 (Robert & Gouet, 2014).

**Figure S4.**
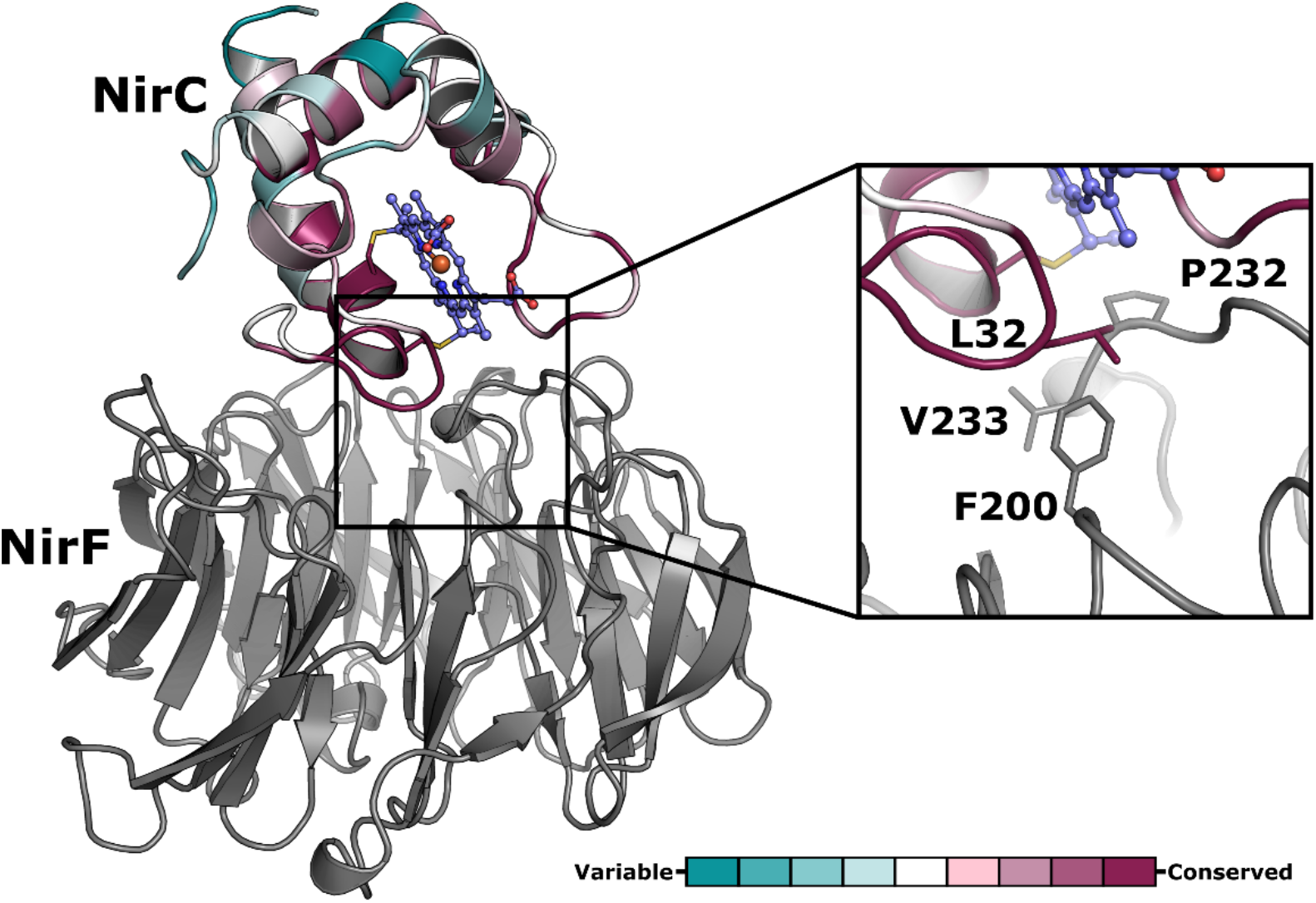
Depiction of NirC (coloured according to sequence conservation) and NirF (grey) (PDB: 6TV2) after superposition onto the cytochrome *c* and *d_1_*-domains of nitrite reductase NirS (PDB: 1nir), respectively.

**Figure S5.**
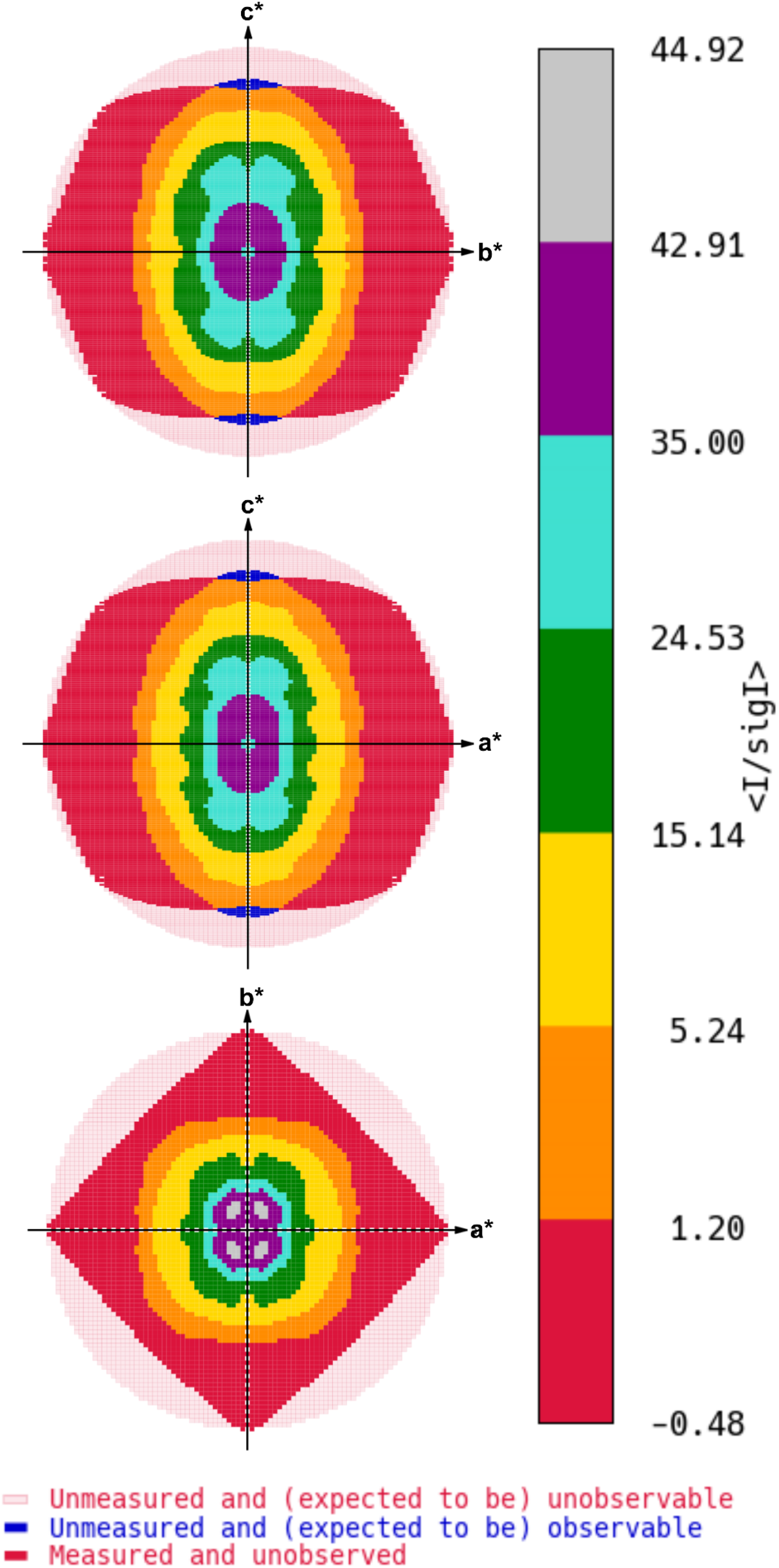
Graphical representation of the <I/σ(I)>-values along the principal axes of the reciprocal lattice.

**Figure S6.**
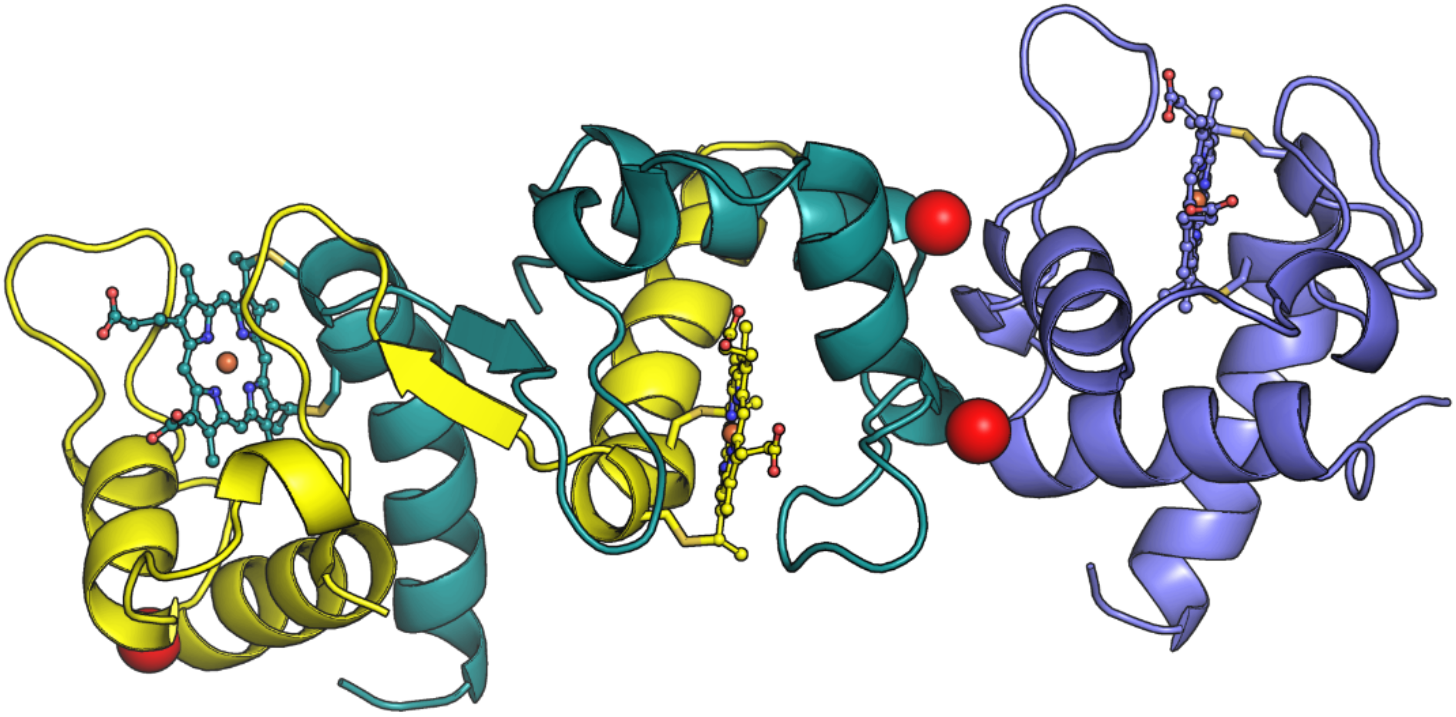
Depiction of chain G (blue), J (teal) and D (yellow) with a red sphere highlighting the mutation site of E71A, which was introduced to increase the crystallisability of NirC.

## References

Afonine, P. V., Grosse-Kunstleve, R. W., Echols, N., Headd, J. J., Moriarty, N. W., Mustyakimov, M., Terwilliger, T. C., Urzhumtsev, A., Zwart, P. H. & Adams, P. D. (2012). Acta crystallographica. Section D, Biological crystallography. 68, 352–367, doi:10.1107/S0907444912001308.

Almagro Armenteros, J. J., Tsirigos, K. D., Sønderby, C. K., Petersen, T. N., Winther, O., Brunak, S., Heijne, G. von & Nielsen, H. (2019). Nature biotechnology. 37, 420–423, doi:10.1038/s41587-019-0036-z.

Altenhoff, A. M., Glover, N. M., Train, C.-M., Kaleb, K., Warwick Vesztrocy, A., Dylus, D., Farias, T. M. de, Zile, K., Stevenson, C., Long, J., Redestig, H., Gonnet, G. H. & Dessimoz, C. (2018). Nucleic acids research. 46, D477–D485, doi:10.1093/nar/gkx1019.

Arai, H., Sanbongi, Y., Igarashi, Y. & Kodama, T. (1990). FEBS Letters. 261, 196–198, doi:10.1016/0014-5793(90)80669-A.

Arslan, E., Schulz, H., Zufferey, R., Künzler, P. & Thöny-Meyer, L. (1998). Biochemical and biophysical research communications. 251, 744–747, doi:10.1006/bbrc.1998.9549.

Ashkenazy, H., Abadi, S., Martz, E., Chay, O., Mayrose, I., Pupko, T. & Ben-Tal, N. (2016). Nucleic acids research. 44, W344–50, doi:10.1093/nar/gkw408.

Bertini, I., Cavallaro, G. & Rosato, A. (2006). Chemical reviews. 106, 90–115, doi:10.1021/cr050241v.

Boer, A. P. de, Reijnders, W. N., Kuenen, J. G., Stouthamer, A. H. & van Spanning, R. J. (1994). Antonie van Leeuwenhoek. 66, 111–127, doi:10.1007/bf00871635.

Brown, K., Nurizzo, D., Besson, S., Shepard, W., Moura, J., Moura, I., Tegoni, M. & Cambillau, C. (1999). Journal of molecular biology. 289, 1017–1028, doi:10.1006/jmbi.1999.2838.

Burkhardt, A., Pakendorf, T., Reime, B., Meyer, J., Fischer, P., Stübe, N., Panneerselvam, S., Lorbeer, O., Stachnik, K., Warmer, M., Rödig, P., Göries, D. & Meents, A. (2016). Eur. Phys. J. Plus. 131, 25, doi:10.1140/epjp/i2016-16056-0.

Cowtan, K. (2006). Acta Crystallographica Section D: Biological Crystallography. 62, 1002–1011, doi:10.1107/S0907444906022116.

Cowtan, K. (2010). Acta crystallographica. Section D, Biological crystallography. 66, 470–478, doi:10.1107/S090744490903947X.

Di Donato, A., Cafaro, V., Romeo, I. & D’Alessio, G. (1995). Protein science: a publication of the Protein Society. 4, 1470–1477, doi:10.1002/pro.5560040804.

Emsley, P. & Cowtan, K. (2004). Acta Crystallographica Section D: Biological Crystallography. 60, 2126–2132, doi:10.1107/S0907444904019158.

Evans, P. R. (2011). Acta crystallographica. Section D, Biological crystallography. 67, 282–292, doi:10.1107/S090744491003982X.

Evans, P. R. & Murshudov, G. N. (2013). Acta crystallographica. Section D, Biological crystallography. 69, 1204–1214, doi:10.1107/S0907444913000061.

Goldschmidt, L., Cooper, D. R., Derewenda, Z. S. & Eisenberg, D. (2007). Protein science: a publication of the Protein Society. 16, 1569–1576, doi:10.1110/ps.072914007.

Hasegawa, N., Arai, H. & Igarashi, Y. (2001). Biochemical and biophysical research communications. 288, 1223–1230, doi:10.1006/bbrc.2001.5919.

Hayashi, Y., Nagao, S., Osuka, H., Komori, H., Higuchi, Y. & Hirota, S. (2012). Biochemistry. 51, 8608–8616, doi:10.1021/bi3011303.

Hirota, S. (2019). Journal of inorganic biochemistry. 194, 170–179, doi:10.1016/j.jinorgbio.2019.03.002.

Hirota, S., Hattori, Y., Nagao, S., Taketa, M., Komori, H., Kamikubo, H., Wang, Z., Takahashi, I., Negi, S., Sugiura, Y., Kataoka, M. & Higuchi, Y. (2010). Proceedings of the National Academy of Sciences of the United States of America. 107, 12854–12859, doi:10.1073/pnas.1001839107.

Holm, L. & Laakso, L. M. (2016). Nucleic acids research. 44, W351–5, doi:10.1093/nar/gkw357.

Hu, X., Wang, H., Ke, H. & Kuhlman, B. (2007). Proceedings of the National Academy of Sciences of the United States of America. 104, 17668–17673, doi:10.1073/pnas.0707977104.

Junedi, S., Yasuhara, K., Nagao, S., Kikuchi, J.-I. & Hirota, S. (2014). Chembiochem: a European journal of chemical biology. 15, 517–521, doi:10.1002/cbic.201300728.

Kabsch, W. (2010). Acta Crystallographica Section D: Biological Crystallography. 66, 125–132, doi:10.1107/S0907444909047337.

Kawasaki, S., Arai, H., Igarashi, Y. & Kodama, T. (1995). Gene. 167, 87–91, doi:10.1016/0378-1119(95)00641-9.

Klünemann, T., Preuß, A., Adamczack, J., Rosa, L. F. M., Harnisch, F., Layer, G. & Blankenfeldt, W. (2019). Journal of molecular biology. 431, 3246–3260, doi:10.1016/j.jmb.2019.05.046.

Koon, N., Squire, C. J. & Baker, E. N. (2004). Proceedings of the National Academy of Sciences of the United States of America. 101, 8295–8300, doi:10.1073/pnas.0400820101.

Korasick, D. A., Westfall, C. S., Lee, S. G., Nanao, M. H., Dumas, R., Hagen, G., Guilfoyle, T. J., Jez, J. M. & Strader, L. C. (2014). Proceedings of the National Academy of Sciences of the United States of America. 111, 5427–5432, doi:10.1073/pnas.1400074111.

Liu, Y. & Eisenberg, D. (2002). Protein science: a publication of the Protein Society. 11, 1285–1299.

Matsuura, Y., Takano, T. & Dickerson, R. E. (1982). Journal of molecular biology. 156, 389–409, doi:10.1016/0022-2836(82)90335-7.

Miroux, B. & Walker, J. E. (1996). Journal of molecular biology. 260, 289–298, doi:10.1006/jmbi.1996.0399.

Murshudov, G. N., Skubák, P., Lebedev, A. A., Pannu, N. S., Steiner, R. A., Nicholls, R. A., Winn, M. D., Long, F. & Vagin, A. A. (2011). Acta Crystallographica Section D: Biological Crystallography. 67, 355–367, doi:10.1107/S0907444911001314.

Nagao, S., Ueda, M., Osuka, H., Komori, H., Kamikubo, H., Kataoka, M., Higuchi, Y. & Hirota, S. (2015). PloS one. 10, e0123653, doi:10.1371/journal.pone.0123653.

Nurizzo, D., Cutruzzolà, F., Arese, M., Bourgeois, D., Brunori, M., Cambillau, C. & Tegoni, M. (1998). Biochemistry. 37, 13987–13996, doi:10.1021/bi981348y.

Parui, P. P., Deshpande, M. S., Nagao, S., Kamikubo, H., Komori, H., Higuchi, Y., Kataoka, M. & Hirota, S. (2013). Biochemistry. 52, 8732–8744, doi:10.1021/bi400986g.

Pearson, I. V., Page, M. D., van Spanning, R. J. M. & Ferguson, S. J. (2003). Journal of bacteriology. 185, 6308–6315, doi:10.1128/jb.185.21.6308-6315.2003.

Ren, C., Nagao, S., Yamanaka, M., Komori, H., Shomura, Y., Higuchi, Y. & Hirota, S. (2015). Molecular bioSystems. 11, 3218–3221, doi:10.1039/c5mb00545k.

Robert, X. & Gouet, P. (2014). Nucleic acids research. 42, W320–4, doi:10.1093/nar/gku316.

Schobert, M. & Jahn, D. (2010). International journal of medical microbiology: IJMM. 300, 549–556, doi:10.1016/j.ijmm.2010.08.007.

Schrödinger, L. L. C. (2015). The PyMOL Molecular Graphics System, Version 1.8.

Sheldrick, G. M. (2008). Acta crystallographica. Section A, Foundations of crystallography. 64, 112–122, doi:10.1107/S0108767307043930.

Sievers, F., Wilm, A., Dineen, D., Gibson, T. J., Karplus, K., Li, W., Lopez, R., McWilliam, H., Remmert, M., Söding, J., Thompson, J. D. & Higgins, D. G. (2011). Molecular systems biology. 7, 539, doi:10.1038/msb.2011.75.

Skubák, P. & Pannu, N. S. (2013). Nature communications. 4, 2777, doi:10.1038/ncomms3777.

Swapna, L. S., Srikeerthana, K. & Srinivasan, N. (2012). PloS one. 7, e36688, doi:10.1371/journal.pone.0036688.

Thorn, A. & Sheldrick, G. M. (2011). Journal of applied crystallography. 44, 1285–1287, doi:10.1107/S0021889811041768.

Tickle, I. J., Flensburg, C., Keller, P., Paciorek, W., Sharff, A., Vonrhein, C. & Bricogne, G. (2018). STARANISO.

Vonrhein, C., Flensburg, C., Keller, P., Sharff, A., Smart, O., Paciorek, W., Womack, T. & Bricogne, G. (2011). Acta crystallographica. Section D, Biological crystallography. 67, 293–302, doi:10.1107/S0907444911007773.

Winn, M. D., Ballard, C. C., Cowtan, K. D., Dodson, E. J., Emsley, P., Evans, P. R., Keegan, R. M., Krissinel, E. B., Leslie, A. G. W., McCoy, A., McNicholas, S. J., Murshudov, G. N., Pannu, N. S., Potterton, E. A., Powell, H. R., Read, R. J., Vagin, A. & Wilson, K. S. (2011). Acta crystallographica. Section D, Biological crystallography. 67, 235–242, doi:10.1107/S0907444910045749.

Winter, G., Waterman, D. G., Parkhurst, J. M., Brewster, A. S., Gildea, R. J., Gerstel, M., Fuentes-Montero, L., Vollmar, M., Michels-Clark, T., Young, I. D., Sauter, N. K. & Evans, G. (2018). Acta Crystallographica. Section D, Structural Biology. 74, 85–97, doi:10.1107/S2059798317017235.

Ye, R. W., Arunakumari, A., Averill, B. A. & Tiedje, J. M. (1992). Journal of bacteriology. 174, 2560–2564.

Zajicek, R. S., Bali, S., Arnold, S., Brindley, A. A., Warren, M. J. & Ferguson, S. J. (2009). The FEBS journal. 276, 6399–6411, doi:10.1111/j.1742-4658.2009.07354.x.

